# RAC1 Regulates Shh-Medulloblastoma Growth via GLI-Mediated Transcription

**DOI:** 10.1101/2025.05.22.655563

**Authors:** Nitish Jangde, Mi Hye Lee, Luz Ruiz, Isabelle Egan, Anna M. Jermakowicz, Daniel Wynn, Erik Goka, Marc Lippman, David J. Robbins, Nagi G. Ayad

## Abstract

Medulloblastoma (MB) is the most common malignant primary pediatric brain tumor. Current therapies are ineffective for targeting proliferation, leptomeningeal migration, and metastasis of MB cancer cells to visceral organs and therefore, novel treatments are needed. The small GTPase, RAC1, has emerged as an important regulator of actin cytoskeletal dynamics, proliferation, and migration in several cancers. However, it has not been characterized in MB and no clinical drug candidates have been described for RAC1 in MB. Here we demonstrate that RAC1 levels are higher in MB tissue relative to normal cerebellum. Further, RAC1 depletion significantly reduces proliferation and migration of Shh-MB cells *in vitro*. Mechanistically, RAC1 controls the mRNA and protein levels of the main transcription factors in the Shh pathway, GLI1 and GLI2. RAC1 binds to the GLI1 promoter highlighting a novel role in transcriptional regulation in Shh-dependent cancers. We demonstrate that the RAC1 inhibitor, GYS32661, is brain penetrant, and reduces MB growth and increases mouse survival in an orthotopic model of Shh-MB. Importantly, GYS32661 is a non-toxic clinical candidate, suggesting that it may be a novel potential drug for the treatment of either the pediatric or adult forms of MB. Collectively, our studies identify RAC1 as a druggable target in Shh-dependent MB.

**Graphical abstract:** **RAC1 controls Shh-MB progression by binding to the GLI1 promoter.**

Graphical abstract demonstrating RAC1 localizes to the nucleus and binds to the GLI1 promoter and plays an important role in its transcription. RAC1 genetic or GYS32661 mediated inhibition leads to RAC1 dissociation from the GLI1 promoter and transcriptional repression of Shh-MB biomarkers GLI2, DNMT1 and UHRF1. This eventually causes inhibition of Shh-MB tumor cell proliferation and migration, which leads to a decrease in Shh-MB development.

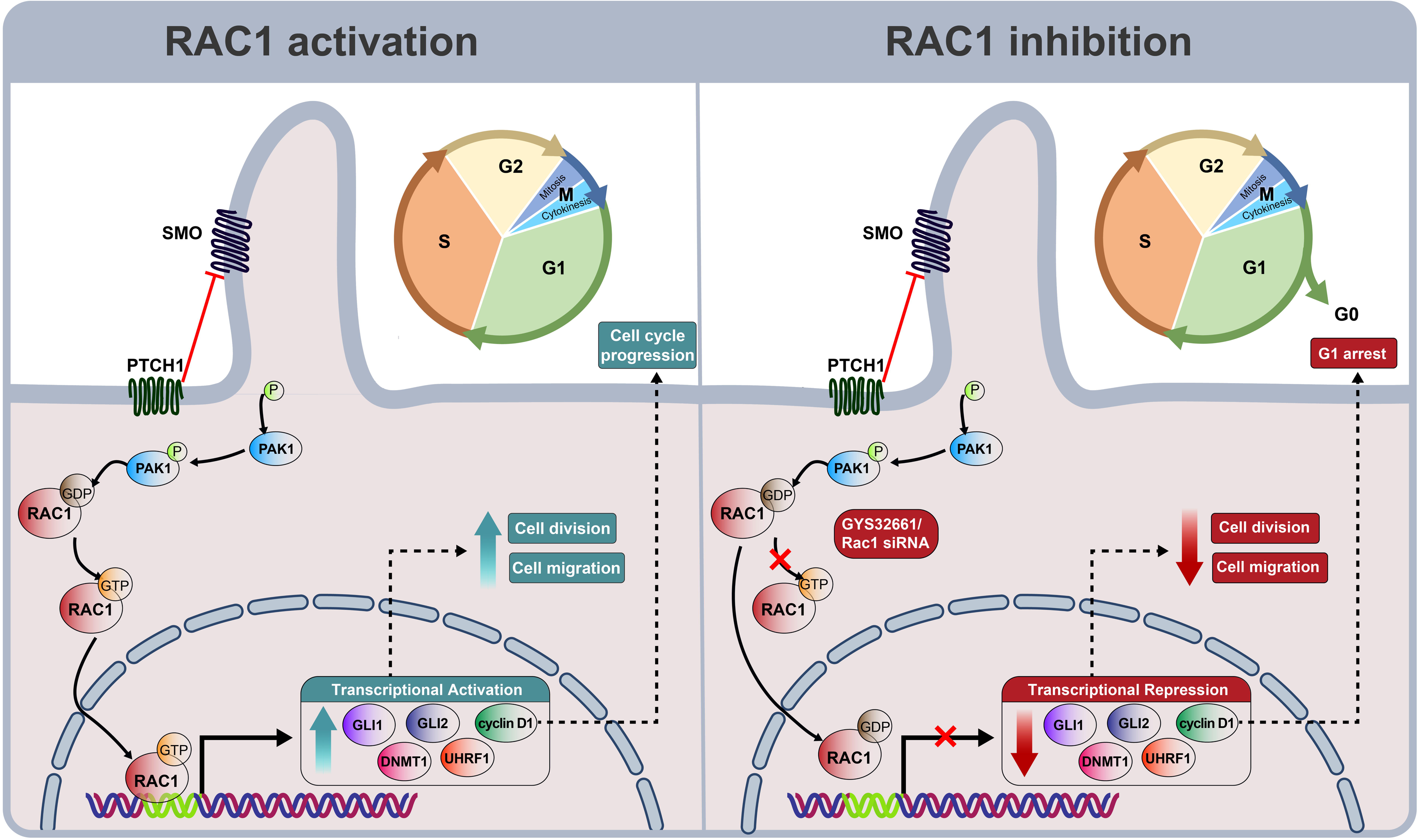

## Introduction

Medulloblastoma (MB) is the most prevalent malignant primary brain tumor in children, and a leading cause of childhood cancer-related illness and death(1). Current treatment strategies involve surgery, radiation (for children older than three years old), and chemotherapy(2). MB is categorized into four distinct molecular subgroups: Wingless (Wnt), Sonic Hedgehog (Shh), Group 3, and Group 4. The Shh subgroup is complex, affecting patients across a wide age range—from infants to adults—and is further subdivided into four subtypes: Shh-α (adolescents), Shh-β (infants with a poor prognosis), Shh-γ (infants with a favorable prognosis), and Shh-δ (adults)(3). It accounts for approximately 30% of all medulloblastomas, and five-year survival is about 70% for Shh-MB patients^30^. Shh-MB most frequently occurs in children under three years of age and above 17 years of age, with fewer cases reported in childhood and adolescence(4). It originates in the cerebellum from cerebellar granule progenitor cells with activation of the Shh pathway, where Shh dysregulation triggers Shh-MB tumor growth during embryonic brain development (5, 6). For example, mutations of receptors in the Shh pathway, PTCH1 and SMO (Smoothened), are often associated with Shh-MB development(7). In terms of therapeutic interventions that have been tried, SMO antagonists as well as inhibitors of the downstream transcriptional regulators of the pathway GLI1 and GLI2(8–11) have been reported. However, resistance has been observed in both preclinical and clinical settings, suggesting the need for novel therapeutic strategies(12–15).

Ras-related C3 botulinum toxin substrate 1 (RAC1) overexpression has been implicated in several cancers, such as breast cancer, non-small cell lung cancer, and head and neck squamous cell carcinomas(16–18). Emerging evidence suggests that RAC1 may be a therapeutic target in cancer due to its involvement in various cellular processes(19–23). Our study uncovers a new mechanism in which RAC1 controls the levels of several mediators of the Shh pathway, including GLI1. RAC1 depletion or inhibition reduces the transcription of *GLI1*, *GLI2*, and various transcriptional and epigenetic regulators involved in Shh signaling. Consistent with this finding, RAC1 inhibition with the brain penetrant molecule GYS32661 reduces medulloblastoma growth *in vivo*. Our study describes the RAC1 pathway as a point of therapeutic intervention for pediatric and adult Shh-MB.

## Results

### RAC1 is overexpressed in medulloblastoma tumors

As RAC1 has not been extensively characterized in MB we wanted to determine its levels in tumors relative to normal cerebellum. Microarray analysis from de Bont et al. 2008(24) revealed that *RAC1* mRNA was significantly elevated in MB compared to normal cerebellum (Figure 1A) Consistent with these findings, analysis of single-cell RNA-sequencing (scRNA-seq) datasets revealed significantly elevated *RAC1* mRNA expression in neoplastic MB cells (Hovestadt et al., 2013(25)) compared to non-neoplastic cells (Darmanis et al. 2015(26)) (Figure 1B). To determine whether the elevated expression was specific to tumor cells, we compared *RAC1* levels across various non-neoplastic cell populations, including oligodendrocytes, microglia, neural, endothelial and astrocytes. Even when accounting for variable expression in non-neoplastic cell populations, *RAC1* expression was still most pronounced in neoplastic cells (Figure 1C). As shown in Figure 1D, RAC1 protein is higher in human MB tissue relative to cerebellum as judged by immunohistochemistry. In addition, RAC1 protein expression was assayed in several human cancer cells, including MB. Among these, the highest expression was observed in ONS76, which are adherent Shh-MB cells (Supplementary Figure 1). Collectively, these studies indicate that RAC1 expression is higher in MB relative to normal cerebellum and that ONS76 cells may be a good model system to probe RAC1 activity *in vitro*.

**Figure 1.**
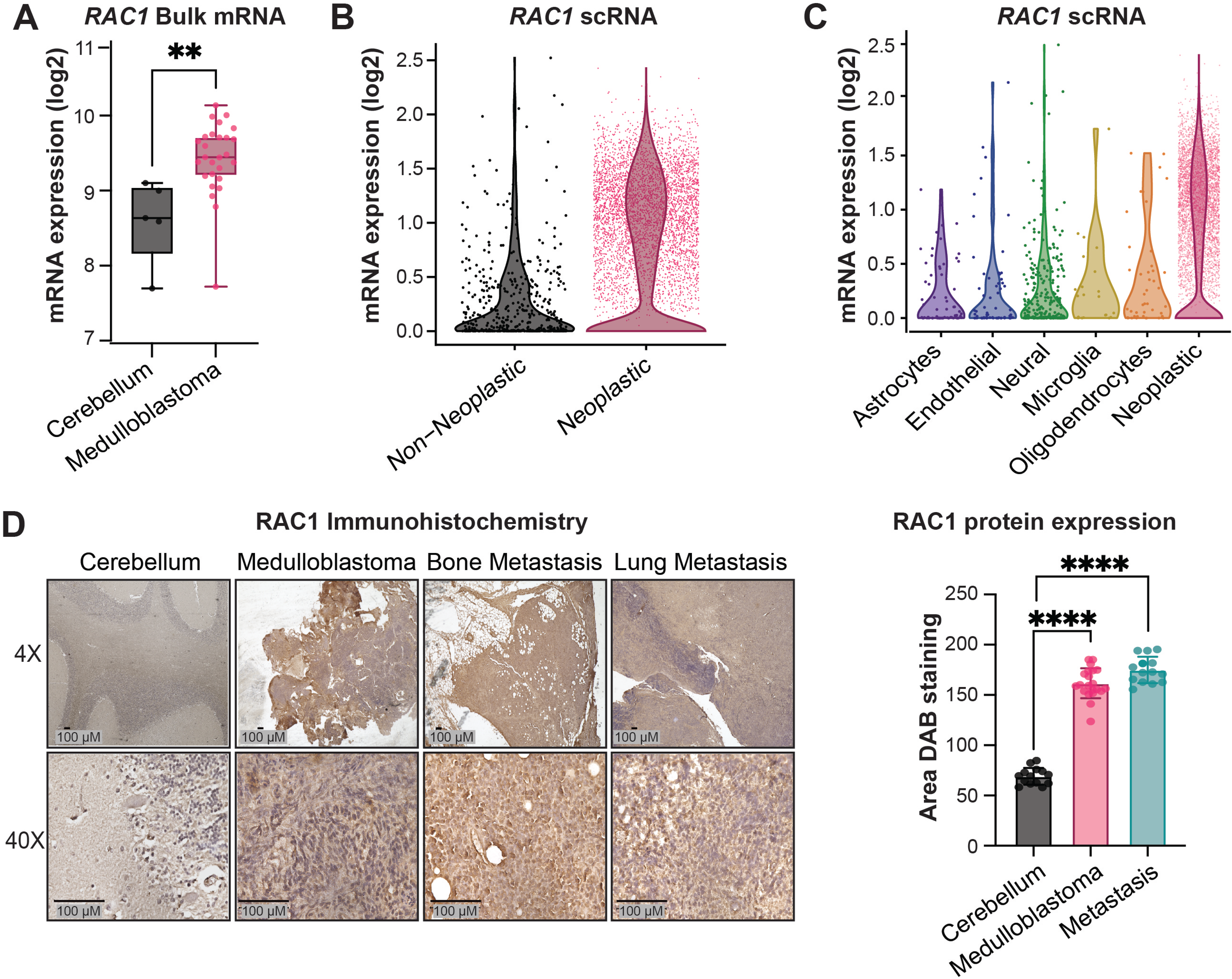
RAC1 is overexpressed in medulloblastoma tumors relative to normal cerebellum. (A) RAC1 mRNA expression in normal cerebellum and medulloblastoma samples from de Bont et al. 2008 microarray data. (B) RAC1 expression levels in neoplastic and non-neoplastic cell populations based on scRNA-seq analysis of MB tumors and healthy brain tissue. (C) RAC1 expression levels in neoplastic cells versus various non-neoplastic cell types, including oligodendrocytes, microglia, neural, endothelial and astrocytes cells. (D) Representative images and quantification of immunohistochemical staining of human cerebellum, MB tumor, MB lung metastasis, and MB bone metastasis for RAC1 (means ±SD, different fields of cerebellum n=2, MB n=3, lung n=1, bone n=1, p values by unpaired 2-tailed t-Test *P<0.05, **P<0.001, ****P<0.0001) (Source data file).

### RAC1 regulates the expression of GLI1 and GLI2 and associated proteins

Since ONS76 cells showed the highest RAC1 expression among all tested cells, we selected them for transfection with either siRNAs targeting *RAC1* or control. As shown in Figure 2A and 2E, we achieved greater than 80% knockdown of RAC1 as measured by real-time qPCR and Western blot analysis and real-time qPCR. Importantly, we observed reduced GLI1 mRNA and protein when RAC1 was depleted (approximately 70%, Figure 2A-B and 2E). As GLI1 is a major transcription factor in the Shh pathway we sought to determine whether other components of the Shh pathway also decreased. Indeed, when we analyzed another transcriptional activator in the Shh pathway, GLI2, it was similarly decreased in *RAC1* siRNA transfected cells relative to control cells (Figure 2C and 2E). Our prior studies described a novel complex of GLI1, GLI2, and the epigenetic regulators DNMT1 and UHRF1(27). Both DNMT1 and UHRF1 are required for Shh signaling and therefore we hypothesized that RAC1 may also control their transcription and protein levels. Indeed, RAC1 siRNA transfected ONS-76 cells had reduced levels of DNMT1 and UHRF1 mRNA and protein (Figure 2D-E). These studies suggest that RAC1 regulates the Shh pathway in MB cells by controlling the expression levels of *GLI1*, *GLI2*, *DNMT1,* and *UHRF1*.

**Figure 2.**
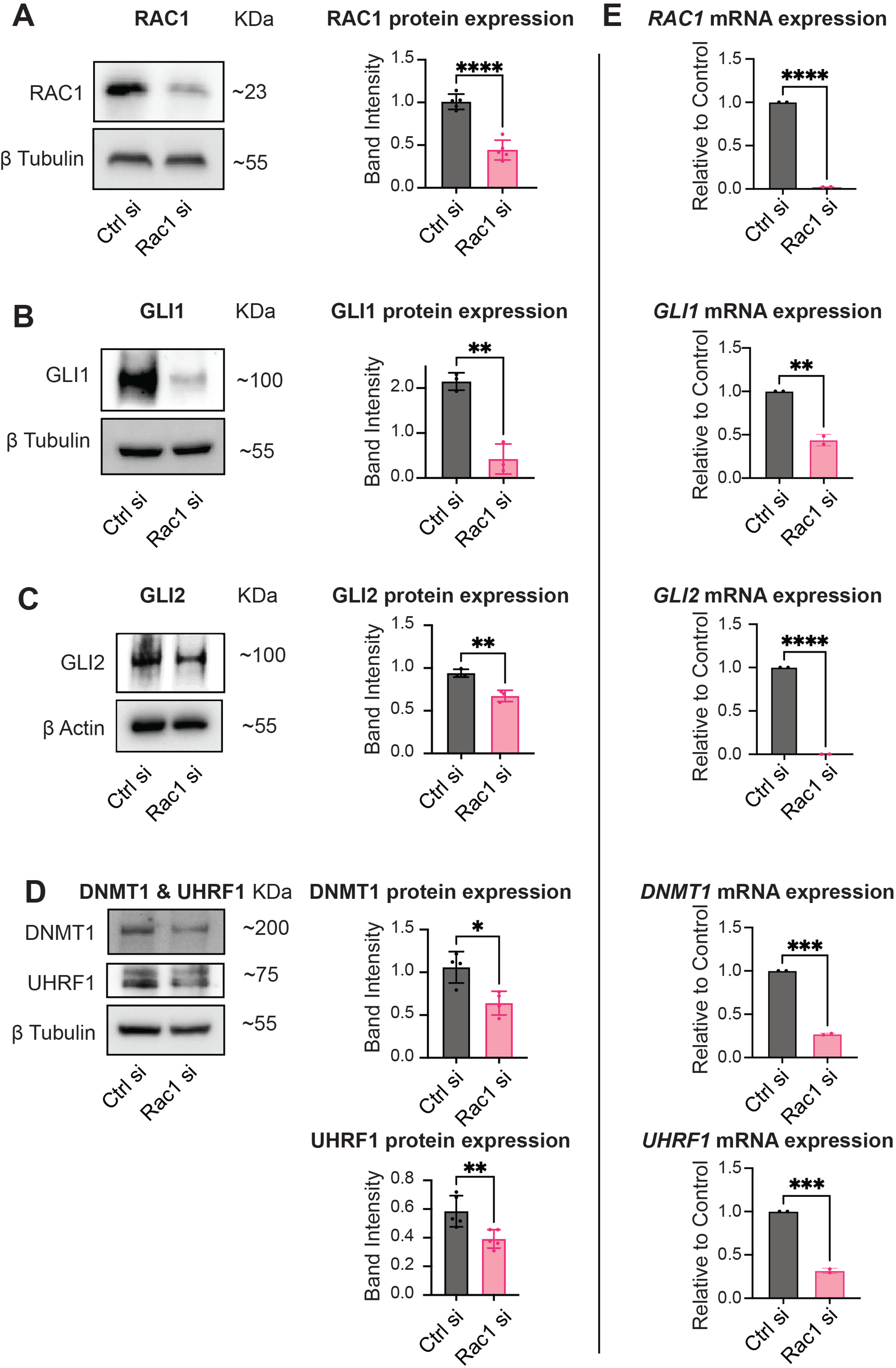
RAC1 depletion in Shh-MB cells reduces levels of GLI1, GLI2, DNMT1 and UHRF1. (A-D) Representative immunoblots for RAC1, GLI1, GLI2, DNMT1 and UHRF1 levels, and their corresponding loading control (β Tubulin or β Actin) after Rac1 siRNA mediated depletion in ONS76 cells; (means ±SD, n= 3-5, p values by unpaired 2-tailed t-Test *P<0.05, **P<0.001, ****P<0.0001), (E) Relative mRNA expression of RAC1, GLI1, GLI2, DNMT1 and UHRF1 in ONS76 cells (means ±SD, n= 2; p values by unpaired 2-tailed t-Test *P<0.05, **P<0.001, ****P<0.0001) (Source data file).

### The RAC1 inhibitor GYS32661 reduces levels of GLI1 and GLI2 *in vitro*

To complement the *RAC1* knockdown studies in Shh-MB, we assessed the effect of pharmacological inhibition of RAC1 in ONS76 cells. We selected the recently reported RAC inhibitor GYS32661(28–31) as our studies demonstrated that it is safe and non-toxic in animals and is brain penetrant (Supplementary Figure S2 and S3). We treated ONS76 cells with varying concentrations of GYS32661 and assessed RAC1 activity (measured as GTP-bound RAC1) using GTP-RAC1 affinity-based purification followed by RAC1 immunoblotting. GYS32661 reduced levels of GTP-RAC1 in ONS76 cells, suggesting that it effectively inhibits RAC1 activity in Shh-MB cells (Figure 3A).

**Figure 3.**
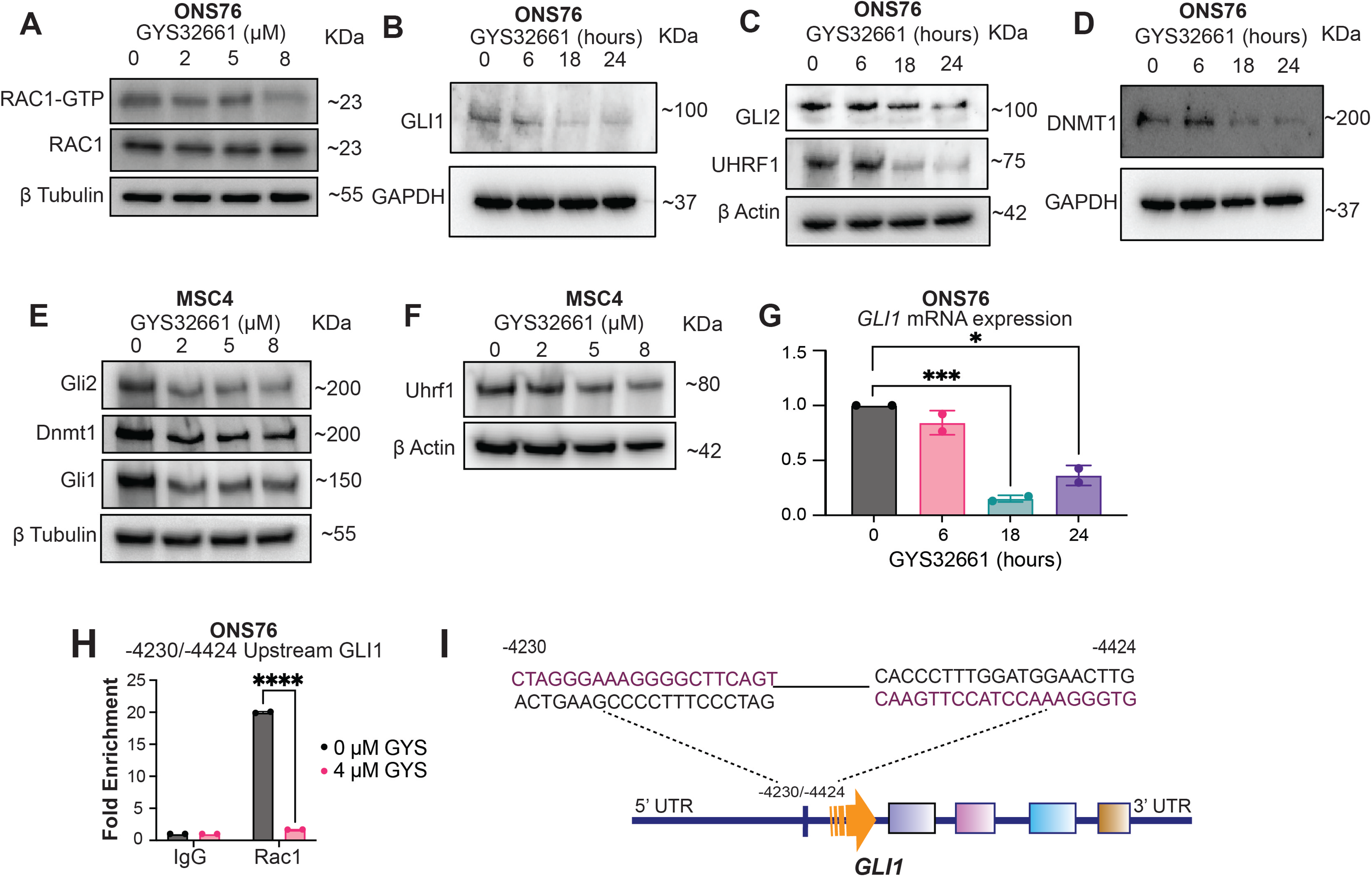
RAC1 inhibition by GYS32661 in Shh-MB cells reduces levels of GLI1, GLI2, DNMT1 and UHRF1. (A) Representative immunoblots showing RAC1-GTP and RAC1 levels with their corresponding loading control (β Tubulin) after treatment with varying GYS32661 concentrations for 24 hours in ONS76 cells. (B) Immunoblot representative images showing GLI1 protein expression in ONS76 cells as compared to its corresponding loading control GAPDH and GLI1, after 8μM, GYS32661 treatment at different time points. (C-D) Immunoblot representative images showing GLI2, UHRF1, and DNMT1 expression as compared to their corresponding loading control (β Actin or GAPDH) in ONS76 cells after GYS32661 treatment at different time points. (E) Representative immunoblots showing GLI1, GLI2 and DNMT1 levels and their corresponding loading control (β Tubulin) after treatment with different concentrations of GYS32661 for 24 hours in MSC4 cells. (F) Representative immunoblots demonstrating UHRF1 levels in MSC4 cells with its corresponding loading control (β Actin) after treatment with different concentrations of GYS32661 for 24 hours. (G) Quantitative representation of relative mRNA levels of GLI1 in ONS76 cells after GYS32661 treatment at different time points (means ±SD, n=2; values by unpaired 2-tailed t-Test *P<0.05, **P<0.001, ****P<0.0001). (H) Quantitative representation of ChIP qPCR of ONS76 cells upon GYS32661 treatment (means ±SD, n=2; values by two-way ANOVA with uncorrected Fisher’s LSD, *P<0.05, **P<0.001, ****P<0.0001 representative of 3 individual replicates); (I) Diagram representing the binding site of RAC1 at the GLI1 upstream promoter region (Source data file).

We next evaluated the downstream effects of RAC1 inhibition on Shh signaling. GYS32661 treatment led to a reduction in GLI1 and GLI2 levels in ONS76 cells (Figure 3B and 3C). Further, GYS32661 reduced Gli1 and Gli2 levels in a model of Shh-MB derived from *Ptch1* knockout mouse tumors (*Ptch1^-/-^*; MSC4 cells), suggesting that this compound can broadly affect Shh-MB cells (Figure 3E and 3F). At the transcriptional level, GYS32661 significantly reduced GLI1 mRNA expression in ONS76 cells (Figure 3G). Consistent with this, GYS32661 treatment decreased mRNA levels of *Gli1, Gli2, Dnmt1* and *Uhrf1* in MSC4 cells (Supplementary Figure S4A).

Prior studies in other cancers suggested that RAC1 can function within a transcriptional complex(32–35), and therefore we wanted to determine whether RAC1 controls *GLI1* levels by binding to chromatin in Shh-MB. To assess this, we performed a Chromatin Immunoprecipitation assay (ChIP) in ONS76 cells to determine whether RAC1 is localized to the *GLI1* promoter. We found that RAC1 binds to a previously characterized *GLI1* promoter region in ONS76 cells (−4230/−4424 *GLI1* upstream, Figure 3I). Among three different regions, the upstream −4230/-4424 showed significantly higher fold enrichment as compared to the IgG control, which suggests that RAC1 binds to this region. Importantly, treatment with GYS32661 for 24 hours reduced RAC1 binding to this region as compared to the DMSO control (Figure 3H-I). Collectively, these studies suggest that RAC1 directly controls *GLI1* transcription by binding to its promoter.

### GYS32661 treatment reduces levels of GLI1-UHRF1-DNMT1 complex *in vitro*

Our prior studies demonstrated that the FDA-approved DNMT1 inhibitor 5-azacytidine, disrupts the GLI-UHRF1-DNMT1 complex, and dramatically reduces Shh-MB growth *in vivo*(27). Therefore, pharmacological targeting of this complex is feasible, which motivated us to test whether GYS32661 would similarly affect the GLI-UHRF1-DNMT1 complex. We treated ONS76 or MSC4 cells with GYS32661 and measured the steady-state levels of the mRNA and protein of all the components of the GLI-UHRF1-DNMT1 complex. As shown in Figure 3A-G, GYS32661 treatment reduced the levels of all components of this complex.

As an independent test, we also checked the interactions of the GLI-UHRF1-DNMT1 complex using an immunofluorescence-based Proximity ligation assay (PLA) (Figure 4A-D). GLI1-DNMT1 interaction was significantly inhibited by GYS32661 compound treatment as the number of PLA puncta per nucleus showing GLI1-DNMT1 interactions were significantly decreased in treated ONS76 and MSC4 cells (Figure 4A-B). Similarly, GYS32661 treatment reduced GLI1-UHRF1 PLA puncta per nuclei in both ONS76 and MSC4 cells (Figure 4C-D). Importantly, siRNA mediated depletion of *RAC1* in ONS76 cells also reduced the GLI1-GLI2 interaction by PLA in ONS76 cells (Supplementary Figure S4C). Altogether, these studies suggest that RAC1 inhibition or depletion reduces the steady-state levels of the components of the GLI-UHRF1-DNMT1 complex, thereby diminishing their interaction in ONS76 and MSC4 cells.

**Figure 4.**
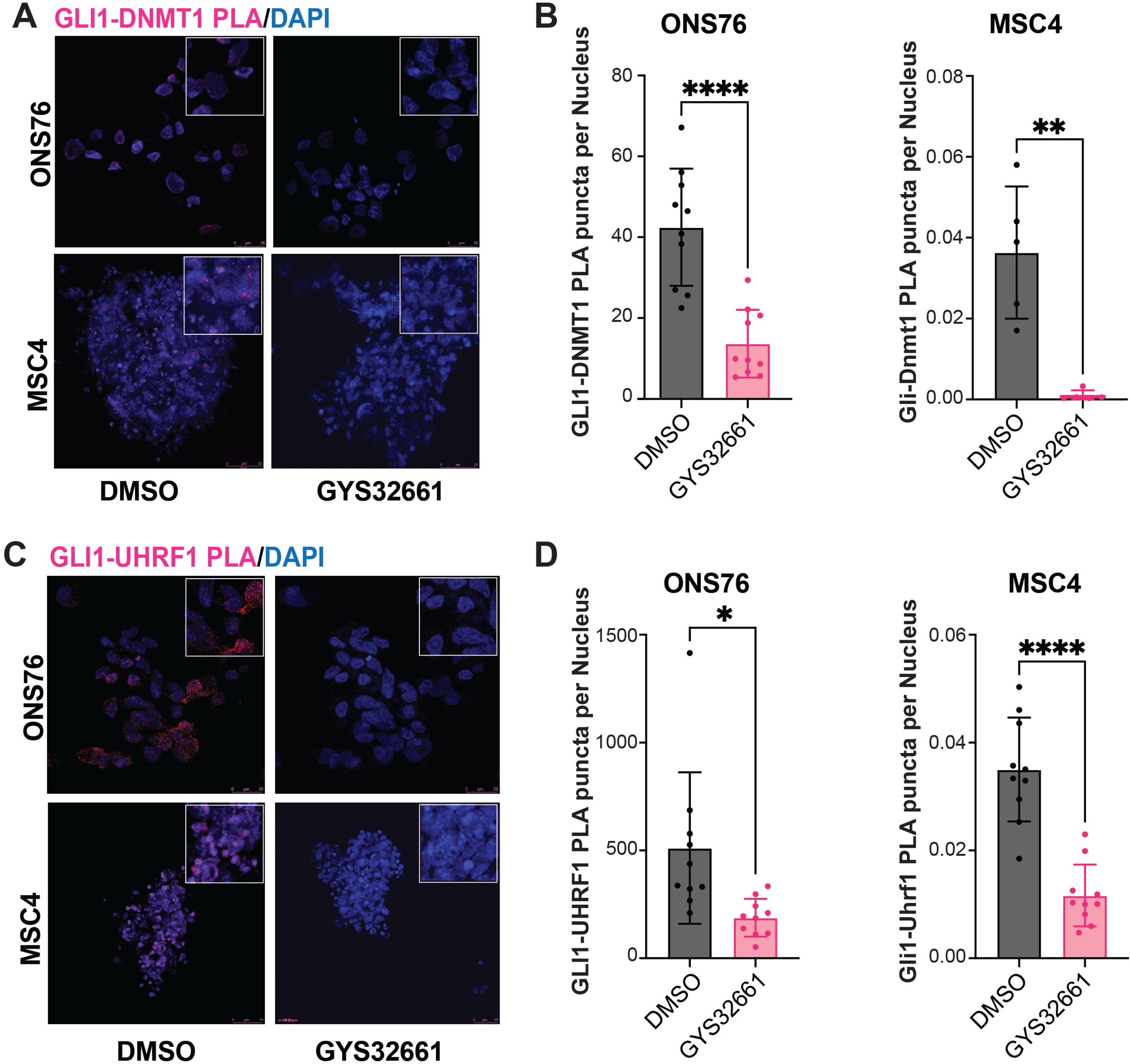
GYS32661 treatment reduces GLI1-GLI2-DNMT1-UHRF1 interactions in Shh-MB cells. (A & B) Representative images of PLA assays of ONS76 and MSC4 cells upon GYS32661 treatment for 24 hours showing the GLI1-DNMT1 interaction, (means ±SD, n= 5-10; values by unpaired 2-tailed t-Test *P<0.05, **P<0.001, ****P<0.0001). (C & D) Representative images of PLA assay of ONS76 and MSC4 cells after GYS32661 treatment for 24 hours showing GLI1-UHRF1 interaction (means ±SD, n= 5-10; values by unpaired 2-tailed t-Test *P<0.05, **P<0.001, ****P<0.0001) (Source data file).

### GYS32661 treatment inhibits cellular motility of ONS76 cells *in vitro*

RAC1 has been shown to have a myriad of roles in proliferation and migration of multiple cell types(20, 21, 36) and therefore we wanted to first test whether RAC1 inhibition would affect Shh-MB cellular migration. We first assessed the efficacy of GYS32661 treatment in a wound healing assay in ONS76 cells. GYS32661 treatment resulted in a marked decrease in cellular migration across the wound, thereby impairing wound healing (Supplementary Figure S5A). A chemotaxis experiment was also performed to assess the efficacy of the GYS32661 compound on cellular movement. ONS76 cells were seeded in very low numbers and movement was tracked by time-lapse imaging in the presence of GYS32661 or vehicle control. Importantly, the path length in GYS32661 treated cells shorter (approximately 280μm) than in control (approximately 380μm), suggesting that this compound inhibited cellular migration in a chemotaxis assay (Supplementary Figure S5B).

We next determined whether disrupting actin dynamics can be part of the reason for reduced migration in GYS32661 cells. Actin polymerization and depolymerization is essential for stress fiber formation, lamellipodia, filopodia, and membrane ruffling, which facilitate cellular motility or migration. The Rho GTPases RhoA, RAC1, and CDC42 are well recognized as critical regulators of actin dynamics and membrane structures (22, 23, 37, 38). To determine the effect of GYS32661 on actin polymerization, ONS76 cells were treated with GYS32661 or DMSO control, then stained with Phalloidin (Alexa Fluor 488; Green) or DAPI (Blue). The results showed a significant decrease in filamentous actin levels after GYS32661 treatment (Supplementary Figure S6A). Interestingly, treated cells had a distinct morphology compared to control cells with less stress fibers and filopodia formation (Supplementary Figure S6A). Therefore, GYS32661 disrupted actin cytoskeleton organization, consistent with the established roles of RAC1 in these processes.

### GYS32661 treatment inhibits cellular proliferation *in vitro*

Multiple studies have shown that proliferation and migration are coupled processes(39), and therefore we wanted to determine whether the effects we observe on migration of ONS76 cells can also be related to proliferation inhibition following GYS32661 treatment. To test this, we treated ONS76 cells with GYS32661 and performed PI-FACS analysis. As seen in Supplementary Figure S6B, GYS32661 treatment arrests ONS76 cells at the G1 phase of the cell cycle after 6 hours, 18 hours, and 24 hours of treatment. This is consistent with the observed reduction in Cyclin D1 levels after treatment (Supplementary Figure S5C and Supplementary Figure S6C), as Cyclin D1 has an established role in G1 progression and RAC1 has previously been implicated in controlling its levels in other cancers(40). These studies suggest that RAC1 controls cell cycle progression of ONS76 cells.

Primary cilia formation is the hallmark of Shh-MB and its disassembly facilitates mitotic division or cellular proliferation(37, 38, 41). We stained GYS32661-treated ONS76 cells for acetylated tubulin and Pericentrin and we found a dramatic increase in acetylated tubulin and Pericentrin positive primary cilia number in treated cells after 24 hours. However, primary cilia length was not affected (Supplementary Figure S7A). Intraciliary transport downstream signaling was also assessed by immunoblotting upon GYS32661 treatment of MSC4 cells. Vav2 and Ift88 phosphorylation was inhibited after GYS32661 treatment of MSC4 cells via dephosphorylation of Pak1 (Supplementary Figure S7B-C). Gsk3β phosphorylation was also inhibited with increasing concentrations of GYS32661 treatment (Supplementary Figure S7C). In addition, acetylated tubulin (AcTub) levels showed a marked increase after 18 and 24 hours of 8 μM of GYS32661 treatment while Arl13b (ADP-ribosylation factor-like protein 13B) was not affected with the treatment (Supplementary Figure S7D). Arl13b is a small GTPase involved in migration and intra-flagellar transport (IFT) signaling(42, 43) while acetylated tubulin is an axoneme marker for primary cilia also involved in intra-flagellar transport. Importantly, tubulin acetylation is associated with reduced proliferation(44). Collectively, these studies suggest that GYS3266’s effects on ONS76 and MSC4 cells may be related to cell cycle progression and signaling.

### GYS32661 reduces Shh-MB growth and is well-tolerated and brain penetrant *in vivo*

Given that GYS32661 treatment reduced proliferation of Shh-MB cells *in vitro*, we wanted to determine whether it could potentially inhibit tumor growth *in vivo*. To test this, we performed mice allograft experiments with MSC4 cells (Figure 5 A-D). Shh-MB tumors started developing after 2 weeks, which we confirmed by MRI. We treated mice in two groups: one with vehicle (normal saline; NaCl in sterile pH 9.0) and another with GYS32661 (25 mg/kg body weight in sterile normal saline). The mice were treated for 3 weeks, and samples were collected after euthanasia and perfusion when the mice showed neurological symptoms. Once mice started showing classical MB neurological symptoms such as head tilting, difficulty in walking, lethargy, or ataxia, their brains were scanned by MRI to detect MB tumors. GYS32661-treated mice showed a marked reduction in Shh-MB tumor development compared to vehicle-treated mice (Figure 5A-C). Notably, mice showed a statistically significant increase in survival upon GYS32661 treatment (Figure 5D).

**Figure 5.**
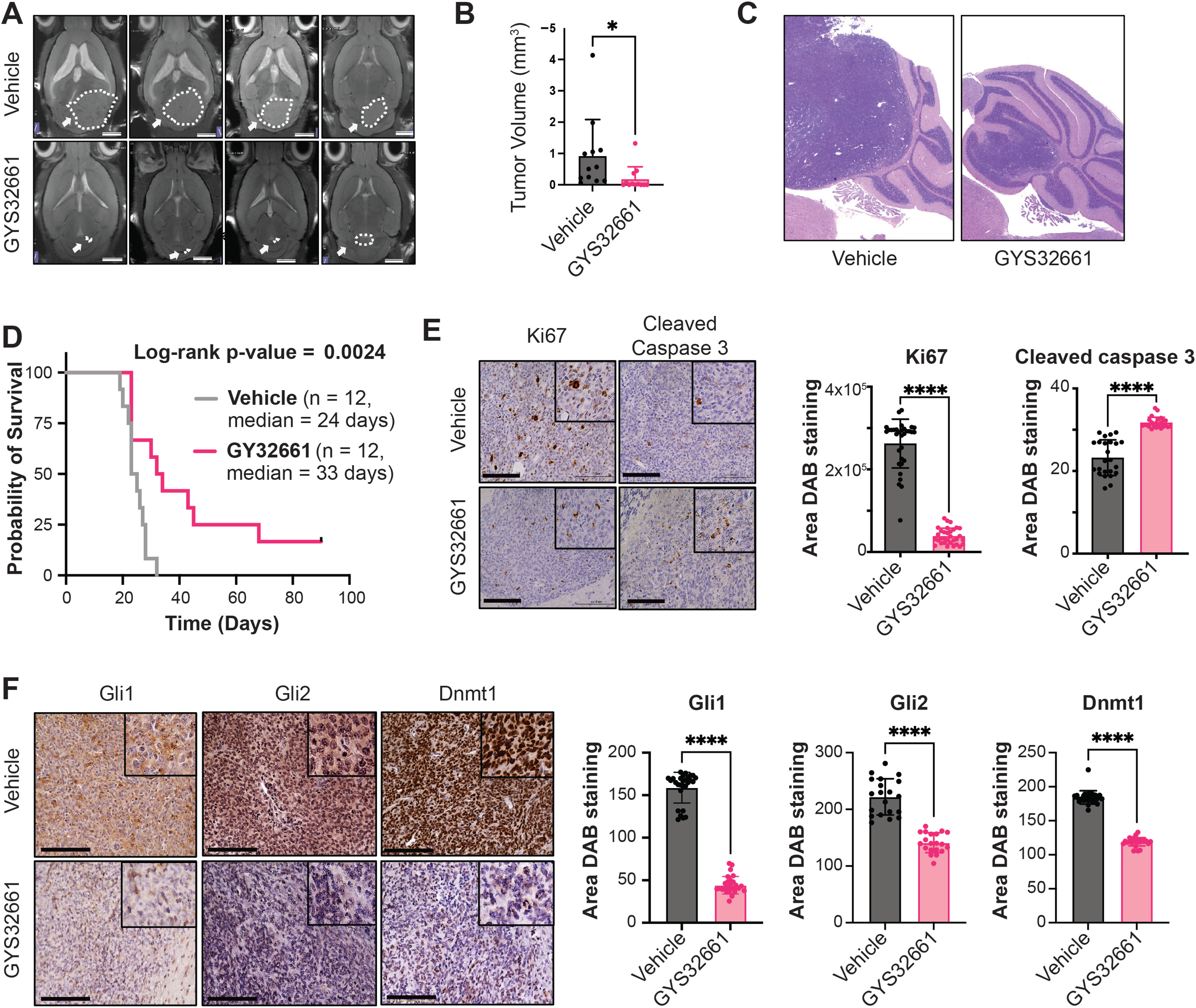
GYS32661 treatment reduces Shh-MB growth *in vivo*. (A & B) Representative MRI images of Shh-MB tumors after *in vivo* MSC4 cell orthotopic injection after endpoint of the vehicle (Upper images in A) or GYS32661 (Lower images in A) treatment, and their representative H&E images, respectively; (B) Quantitation of tumor volumes of orthotopic MSC4 mouse Shh-MB tumors after the endpoint of GYS32661 cohort (means ±SD, n= 12 individual mice per group; values by unpaired 2-tailed t-Test *P<0.05, **P<0.001, ****P<0.0001); (D) Kaplan Meier survival curve analysis of orthotopic MSC4 mouse Shh-MB tumors of GYS32661 cohort (means ±SD, n= 12 individual mice per group). (E & F) Representative IHC images of MSC4 mouse orthotopic Shh-MB tumors after the GYS32661 cohort demonstrating Ki67, Cleaved Caspase-3, GLI1, GLI2 and DNMT1 protein levels, respectively; (means ±SD, n=7 different fields of n=4 representative individual mice; values by unpaired 2-tailed t-Test *P<0.05, **P<0.001, ****P<0.0001) (Source data file).

Tumor tissues were collected at 3 weeks, fixed in 4% formalin and subsequently embedded in paraffin. These paraffin-embedded tumor samples were sectioned and stained for histological markers for proliferation (Ki67) and cell death or apoptosis markers (cleaved caspase-3). GYS32661-treated tumors showed markedly reduced Ki67 staining compared to vehicle-treated mice MB tumors and had higher apoptosis as judged by cleaved caspase-3 (Figure 5E). GYS32661 treatment also reduced levels of on-target biomarkers in the Shh pathway such as GLI1 and GLI2 *in vivo*, suggesting that the compound reaches the tumor site (Figure 5F). Collectively, these studies suggest that GYS32661 increases apoptosis while reducing proliferation and Shh signaling *in vivo*.

In addition to effectively reducing Shh-MB tumor burden *in vivo*, GYS32661 was well-tolerated in animals. Tumor bearing mice treated with GYS32661 did not show any reduction in body weight compared to untreated mice (Supplementary Figure S2A). Similarly, toxicological studies in rats suggest that GYS32661 has no negative effect on the body weight of either male or female rats (Supplementary Figures S2B-C). Further, the dose used in our efficacy studies (25mg/kg) is well-below the maximum tolerated dose tested in rats (45 mg/kg; Supplementary Figure S2B-C). To assess the effect of treatment on the visceral organs, liver and brain weights were also measured in the same groups of male and female rats and no observable toxicity was found (0-45 mg/kg/day for 28 days; Supplementary Figures S3A-D). Importantly, the compound is 50% brain penetrant as determined by HPLC analysis of brain and plasma levels (Supplementary Figures S3E-F). Collectively, these studies suggest that GYS32661 is brain penetrant and safe in animal models.

## Discussion

Our present study provides compelling evidence that RAC1 controls Shh-MB growth by regulating GLI1 and GLI2 levels. Genetic or pharmacological inhibition of RAC1 reduced GLI1 and GLI2 levels *in vitro*. In addition, GYS32661 reduced tumor growth and increased survival in a mouse model of Shh-MB. It is also safe in multiple animal models. Collectively, our findings demonstrate that GYS32661 is a promising treatment for Shh-MB.

Our studies suggest that there is a likely therapeutic window 35mg/Kg body weight or less for GYS32661 since RAC1 is expressed higher in Shh-MB tumors relative to normal brain. Microarray and scRNA-seq analysis of neoplastic relative to non-neoplastic cells showed high *RAC1* expression in neoplastic cells, with low expression across various non-neoplastic cell types (Figure 1A-C). Further, RAC1 protein levels were higher in medulloblastoma patient samples relative to normal cerebellum. Notably, RAC1 levels are further increased in metastatic MB compared to non-metastatic MB, suggesting a potential association with tumorigenesis and dissemination (Figure 1D). Collectively, our results suggest RAC1 mRNA and protein levels are higher in neoplastic cells within medulloblastoma tumors relative to normal cerebellum. These findings support the notion that RAC1 plays a role in MB tumorigenesis with limited effect on non-tumor brain tissue, and therefore it may represent a viable therapeutic target for Shh-MB.

Although RAC1 has been described within the context of Shh signaling, we present a new model of how it affects this important pathway in Shh-MB. Prior studies focused on a possible role for RAC1 in GLI1 shuttling from the cytoplasm to the nucleus(32, 33, 35, 45). However, we did not observe any concentration or time point where GYS32661 treatment affected GLI1 shuttling. Rather, under all conditions, we observed a marked reduction in the total mRNA and protein levels of GLI1, GLI2, DNMT1, and UHRF1. This is consistent with a role for RAC1 in transcription of these genes, supported by our findings that RAC1 localizes to the *GLI1* promoter. Therefore, RAC1 may have a similar role in other cancers where it acts in the nucleus to affect transcription of essential genes involved in proliferation, migration, and metastasis.

Given that RAC1 plays multiple roles in different cells it is not surprising that it affects both proliferation and migration of Shh-MB cells. Although it is not possible to separate the effects on migration from proliferation in dividing cells, we favor a model where RAC1 plays a role in proliferation of Shh-MB cells. As shown in our FACS analysis, RAC1 inhibition via GYS32661 treatment increased the percentage of cells in the G1 phase of the cell cycle, reduced G2 and S phase cells, and Cyclin D1 levels (Supplementary Figure S5C and Supplementary Figure S6B-C). This suggests that RAC1 inhibition regulated cell cycle progression, which is consistent with its role in different other cancers(33, 46). Our data suggest that RAC1’s effects on the cell cycle are at least in part due to RAC1 binding to the *GLI1* promoter and affecting *GLI1* transcription, although we acknowledge it is difficult to separate its multiple roles in the cell cycle.

The effects of RAC1 on proliferation can also be related to primary cilia dysfunction, as suggested by other studies(19, 47–49). However, our analysis of GYS32661-treated cells did not indicate a change in the length of primary cilia (Supplementary File S6A). We further noted that treatment with GYS32661 increased α-tubulin acetylation substantially, which is associated with changes in proliferation based on prior studies. By contrast, GYS32661 did not affect Arl13b accumulation, which is associated with longer cilia in other studies(50). Therefore, the effect of GYS32661 on primary cilia number is likely related to its ability to inhibit the cell cycle in ONS76 cells.

Our studies suggest that targeting Shh-MB via inhibiting RAC1 with GYS32661 is a novel means of downregulating GLI1, GLI2, UHRF1, and DNMT1 activity. This indicates GLI proteins as potential druggable targets to treat medulloblastoma patients. Recently, an epigenetic regulatory complex, GLI-UHRF1-DNMT1, has been found to be a key regulator and druggable target of Shh-MB(27). In a genome-wide CRISPR-Cas9 knockout screening of human (DAOY) and murine (SMB21) Shh-MB cells, DNMT1 has been identified as an epigenetic regulator and a potential target for monotherapy(51). Normal cerebellar development and Shh-MB expansion depend on DNMT1. Targeting Shh signaling downstream of SMO, either through its pharmacological inhibition alone or in conjunction with SMO inhibitors, effectively restricts tumor development and increases survival in Shh-MB models(52–54).

UHRF1 is a key regulator of DNA methylation in mammalian cells. Research has also indicated that UHRF1 contributes to the development of tumors (55–57). In a recent study, immunostaining and western blotting analysis of sixty-eight MB patients showed that UHRF1 expression is present in MB tissues but absent in the normal cerebellum(57). In the same study RNAi-mediated UHRF1 downregulation inhibited MB cell proliferation and colony formation, induced G1/G2 cell cycle arrest, elevated p16 expression, and altered CDK4 location. Altogether these results suggest that UHRF1 may be a useful biomarker and treatment target for MB and therefore GYS32661’s ability to decrease UHRF1 levels in Shh-MB tumors is therapeutically advantageous.

Our present study identifies the crosstalk between RAC1-mediated GLI1 activity and the GLI epigenetic complex-mediated GLI1 activity in Shh-MB development. This is the first report of RAC1 inhibition targeting every component of the epigenetic complex (GLI1, GLI2, DNMT1 and UHRF1 complex), suggesting a novel mechanism in RAC1-mediated Shh-MB development. Collectively, our study suggests a novel role for RAC1 in targeting the druggable GLI epigenetic complex by regulating the transcription of the components of the complex, which eventually can restrict Shh-MB development (Graphical abstract) GYS32661’s ability to reduce on-target biomarkers in the Shh pathway in orthotopic models is likely related to its considerable brain penetrance (Supplementary File S3E-F). Although other RAC1 inhibitors have been described (30, 58) to our knowledge this is the first time one has been demonstrated to be brain penetrant and effective in animal models of Shh-MB. These studies and other drug-like characteristics of GYS32661 suggest that a clinical trial should be considered for patients suffering from Shh-MB.

## METHODS

### Data Analysis

Medulloblastoma scRNA-seq data (neoplastic) was obtained from the Hovestadt 2013 and non-tumor brain cell (non-neoplastic) scRNA-seq data was obtained from the Darmanis 2015 study(25, 26) Analysis and visualization of gene expression was performed using the Seurat R package(59–63).Differences in gene expression between neoplastic and non-neoplastic cells was performed using MAST (implemented in Seurat)(64). The de Bont et al 2008(24) microarray data set was acquired from Gliovis (http://gliovis.bioinfo.cnio.es/) and analyzed to compare gene expression of medulloblastoma tumor to normal cerebellum(65).

### Cell culture

Human Shh medulloblastoma cell line ONS76 was grown in RPMI 1640 medium (with ATCC modification) with 10% Fetal bovine Serum (FBS) and 1x Anti-anti at 37°C, 5% CO_2_. Mouse medulloblastoma spheroid cells (MSC4) isolated from Shh MB tumors of genetic conditional *Ptch1* knockout mice were grown in DMEM F12 medium with 1x B27 supplement at 37°C, 5% CO_2_. Ptch1LoxP (B6;129T2-Ptch1tm1Bjw/ WreyJ), and Gfap-Cre (FVB-Tg(GFAP-cre)25Mes/J) strains were obtained from the Jackson Laboratory and bred to generate Ptch1-Gfap mice(66). Cells from spontaneous tumors in these mice were grown in Dulbecco’s modified Eagle’s medium (DMEM)/F12 (Gibco), B27, and penicillin-streptomycin to generate vismodegib-sensitive SHH-MB cultures: SHH-S47 and Ptch1-Gfap. SHH-S47 and Ptch1-Gfap cells were gifted by Dr. Jezebel Rodriguez-Blanco’s laboratory.

### Gene silencing experiment

Cells were transfected with targeting Rac1 siRNA (SantaCruz Biotechnology; Catalog no.: sc-36351) and non-targeting siRNAs (Santa Cruz Biotechnology; Catalog no.: sc-370071)), transfection reagent and transfection media per the manufacturer’s protocol. After knockdown immunoblotting and qPCR experiments were performed for confirmation.

### Cell lysis and immunoblotting

Lysis buffer (Cell Signaling) containing protease and phosphatase inhibitors was added to cells. The resulting extracts were centrifuged at 16,000 x g for 10 minutes at 4 °C to collect the soluble fraction. Protein concentrations were measured by the Qubit Protein Assay (Invitrogen). Subsequently, 40 μg protein of each sample was resolved on SDS-PAGE, transferred on a nitrocellulose membrane, and immunoblotting was performed using respective antibodies.

### RNA isolation and Quantitative Realtime PCR (qRT-PCR)

Total RNA was purified and using the AllPrep DNA/RNA Mini Kit (Qiagen). Reverse transcription was conducted using the SuperScript III First-Strand Synthesis SuperMix (ThermoFisher). Quantitative real-time PCR (qRT-PCR) was conducted with the SsoAdvanced Universal SYBR Green Supermix (Bio-Rad) and gene-specific primers (Supplementary Table S1) on a CFX Opus96 Real-Time qPCR Detection System (Bio-Rad). The ΔΔCt method was used with *Gapdh* or *18S* rRNA as the reference (Supplementary Table S1) to calculate the fold change in gene expression.

### Chromatin immunoprecipitation (ChIP)

ONS76 cells were cross-linked with formaldehyde, lysed, and sonicated. Chromatin was immunoprecipitated with the anti-RAC1 antibody (clone 23A8, Catalog No. 05-389) or a negative control normal Rabbit IgG antibody (Sigma, Catalog No. 12-370). The DNA was recovered after removal of DNA–protein cross-links and subjected to quantitative amplification of the primers specific to the human Gli1 locus(67) (see Supplementary Table S2).

### Wound healing assay

The migratory response of cells to GYS32661 was examined in a scratch assay. Cells were seeded into 12-well plates (3 x 10⁵ cells/well) and allowed to grow to 90–100% confluence, then serum-starved overnight. A scratch was created using a 200 μl sterile pipette tip, and PBS was used to remove cell debris. Cells were then treated with DMSO control, media with 5% FBS (positive control) and media with 2 μM GYS32661 for 18 hours at 37 °C. Cell migration was observed using a Keyence phase contrast microscope BZ-X800, and wound closure was quantified compared to untreated controls by using Fiji wound healing quantification software.

### Chemotaxis assay

The migratory response of cells to GYS32661was examined. Cells were seeded into 8-well chambered slides (Ibidi) in 1 × 10^3^ cells/well density and allowed them to adhere, then serum-starved overnight. Cells were then treated with DMSO control, media, or with 2 μM GYS32661 for 12 hours at 37 °C in live cell imaging incubator chamber. Cellular movement was documented by time lapse imaging on a Keyence phase BZ-X800 contrast microscope, and quantified by Keyence software BZ-X800.

### GTP-bound RAC1 detection

Cells were treated with varying concentrations of the GYS32661 compound. Cells were harvested under non-denaturing conditions by rinsing them with PBS and lysing them with an ice-cold lysis solution (25mM Tris-HCl, pH 7.2, 150mM NaCl, 5mM MgCl_2_, 1% NP-40, and 5% glycerol containing PMSF), followed by incubation on ice. Cell lysates were mixed with EDTA, GTPγS (positive control), or GDP (negative control) and incubated at 30°C. The reaction was terminated with 1 M MgCl₂ (final working concentration 60 mM) followed by cooling on ice for GTPγS or GDP treatment. Subsequently, the treated lysates were subjected to affinity precipitation using glutathione resin and GST-PAK1-PBD. After gentle shaking at 4°C, the resin was thoroughly washed with lysis buffer, and the GTP-bound RAC1 protein was eluted using a reducing sample buffer containing DTT. The samples were then electrophoresed and immunoblotted for RAC1.

### Proximity ligation Assay (PLA)

PLA was performed on cells using a Duolink Proximity Ligation Assay (Millipore Sigma) as described by the manufacturer. The following antibodies were used for PLA: UHRF1 (catalogue no. sc-373750, Santa Cruz Biotechnology, RRID: AB_10947236), DNMT1 (catalogue no. 5032, Cell Signaling Technology, RRID: AB_10548197), GLI1 (catalog no. AF3455, R&D Systems, RRID: AB_2247710), GLI2 (catalog no. AF3635, R&D Systems, RRID: AB_2111902), rabbit IgG (catalog no. 12370, Millipore Sigma). DAPI staining was performed using ProLong Gold Antifade Mountant with DAPI (Invitrogen).

### Immunofluorescence

GLI1/Phalloidin immunofluorescence studies were performed on ONS76 cells after GYS32661 or DMSO treatment. Spheroid cells (MSC4) were dissociated by Accutase (Gibco) treatment for 5 minutes, centrifuged at 2,000 rpm for 5 minutes, and seeded at 5 × 10^5^ density on poly-L-Lysine pre-coated coverslips (Corning) in a 24-well plate.

Cells were fixed with 4% PFA and permeabilized using 0.1% Triton X-100 in PBS for 15 minutes. 1% BSA in PBS was used to block for 45 minutes at room temperature. Slides were then incubated with GLI1 antibody and fluorescent phalloidin staining solution. Confocal LEICA SP8 microscopy was used to take images of cells after washing.

For cilia staining, cilia were counted in 4% PFA fixed cells after staining with Acetylated tubulin (Sigma cat # T6793) and pericentrin (Novus Cat # NB100-61071) antibodies to mark cilia and the centrosome. Acetylated tubulin-positive cilia were counted by Zeiss confocal microscopy (LMS 800) and the % ciliated cells were normalized by Dapi counting.

### In vivo drug dosing

MSC4 spheroid cells were dissociated by passing through the 8μm cell strainer followed by Accutase (Gibco) treatment. Viable cells were counted using a Cellometer followed by Acridine orange/Propidium Iodide (AO/PI) staining. Orthotopically 1 × 10^5^ cells/mouse were injected into the cerebellum. Before injection a 0.25-inch incision was created on the back of the scalp with a sterilized scalpel. A dental drill was utilized to create a implanted into the cerebellum using the coordinates 2 mm down lambda, 2 mm right of the midline suture, and 2 mm deep(66). The mouse was secured on a stereotactic frame by mounting its incisors onto the frame hold. A 10 μL bevel-tipped syringe, which had been preloaded with 4 μL of tumor cell solution, was slowly inserted into the burr hole. After the bevel of the syringe needle went beneath the skull surface, it was moved an additional 3 mm down before being moved up 0.5 mm. The suspension of the tumor cells was slowly injected under continuous pressure over 30 seconds into the cerebellum. After the injection, the needle was retained in position for two minutes before both the needle and syringe were withdrawn slowly. Finally, the wound was closed using an autoclip. The mice were injected intraperitoneally every other day with GYS32661 compound (35 mg/ Kg of body weight) in 0.9% normal saline pH 5.5 and vehicle (0.9% normal saline pH 5.5). MRI imaging was performed to monitor tumor development every week. GYS32661 compound was also administered in different doses to male and female rats as indicated. After 28 days the body, liver and brain weight of the treated rats were recorded. The brain and blood plasma concentrations of GYS32661 were also measured by HPLC.

### Immunohistochemistry (IHC)

The tissue samples were fixed in 10% Formalin in PBS and paraffin embedded. The formalin fixed paraffin embedded (FFPE) tissues were sectioned by microtome up to 4 μm of thickness. Further, for immunohistochemical staining, endogenous peroxidases were blocked in 3% hydrogen peroxide in methanol. These sections were probed with primary antibody RAC1 Mouse Monoclonal antibody (Proteintech; Cat No. 66122-1-Ig), Ki-67 (Ab15580), Cleaved Caspase-3 (Asp175) Antibody (Cell signaling technology; Cat No. 9661), GLI1 Antibody (A-7) (SantaCruz Biotechnology; Cat No. sc-515781), GLI-2 Antibody (R & D Systems; Cat No. AF3635) and DNMT1 (D63A6) XP Rabbit mAb (Cell signaling technology; Cat No. 5032) Tumor tissue sections slides were subsequently incubated with VECTASTAIN® ABC-HRP Kit, Peroxidase (Mouse IgG) (PK-4002) or VECTASTAIN® ABC-HRP Kit, Peroxidase (Rabbit IgG) (PK-4001) secondary antibody. Staining was visualized DAB Substrate Kit, Peroxidase (HRP), with Nickel, (3,3’-diaminobenzidine) (SK-4100) chromogen and counterstained with hematoxylin. The IHC DAB stained images intensity were quantified as area by using ImageJ software.

## Acknowledgements

We acknowledge R21NS135506, R01NS118023, and the BellRinger at Lombardi Comprehensive Cancer Center of Georgetown University for funding. We also acknowledge Ms. Poorna Prakash for her intensive efforts in helping with primary cilia number counting and length measurement. We thank all members of the Ayad laboratory for helpful discussions. We are grateful to Takele Yazew, our laboratory manager, for helping us during animal surgeries. We acknowledge the Shared Resource Facilities at Georgetown University including the HTSR and FACS cores.

## Contribution

NJ designed and performed experiments and wrote and edited manuscript. M-H L performed experiments and edited manuscript. LR performed analysis and edited manuscript. IE performed experiments and edited manuscript. AMJ assisted with data interpretation and generated figures. ML, DW, DR, edited manuscript. NGA designed experiments and edited manuscript.

---

**Supplementary Figure S1. ONS76 expresses higher RAC1 than other cancer cells**. Immunoblots showing RAC1 expression in different cancer cells with their corresponding loading control, and quantification (means ±SD, n= 3). Immunoblots shown are representative of 3 individual biological replicates (Source data file).

**Supplementary Figure S2. GYS32661 is well-tolerated *in vivo* in mice**. (A) Body weight of C57BL/6 mice with intracranial MSC4 cells after GYS32661 treatment. GYS32661 was administered at 35 mg/kg and animals were weighed on days 4, 6, 8, 19, and 12 (means ±SD, n=12 per group). (B-C) Weights of male and female rats after GYS32661 treatment at varying doses for 28 days (means ±SD, n= 20 per group) (Source data file).

**Supplementary Figure S3. GYS32661 is non-toxic and brain penetrant in rats**. (A-D) Male and female rats received intraperitoneal GYS32661 administration at different doses and organs were harvested after 28 days. Liver and brain weights of male or female were collected after for 28 days (means ± SD, n=20 per group). (E-F) Concentration of GYS32661 in the brain (E) and brain to plasma ratio (F) was measured in mice after 10 mg/kg intraperitoneal injection. Measurements were taken at various time intervals post administration (means ± SD, n=2) (Source data file).

**Supplementary Figure S4. GYS32661 treatment reduces transcription of GLI1, components of GLI-UHRF1-DNMT1 complex, and GLI1-GLI2 interaction**. (A) Quantitation of the relative mRNA expression of *Gli1*, *Gli2*, *Dnmt1* and *Uhrf1* in MSC4 cells (means ±SD, n=2; p values by unpaired 2-tailed t-Test *P<0.05, **P<0.001, ****P<0.0001). (B) Representative images of ONS76 cells GLI1-GLI2 PLA assay after control and RAC1 siRNA treatment (GLI1-GLI2 PLA:Red; DAPI: Blue) and its quantitative representation (means ±SD, n=5; p values by unpaired 2-tailed t-Test *P<0.05, **P<0.001, ****P<0.0001) (Source data file).

**Supplementary Figure S5. GYS32661 treatment reduces Shh-MB cell migration *in vitro***. (A) Representative images and quantification of % migration of cells *in vitro* wound healing assay of ONS76 cells to test GYS32661 compound (means ±SD, n=10 different areas; values by 2-tailed Student’s t-Test P<0.05, **P<0.001, ****P<0.0001 representative of 3 individual biological replicates). (B) Quantitation of accumulated distances traveled by ONS76 cells after GYS32661 treatment in a chemotaxis time lapse imaging experiment (means ±SD, n=60 individual cells; values by 2-tailed Student’s t-Test *P<0.05, **P<0.001, ****P<0.0001) (Source data file).

**Supplementary Figure S6. RAC1 inhibition by GYS32661 in Shh-MB cells induces morphological changes and cell cycle arrest.** (A) Representative images of actin polymerization assay of ONS76 cells (Phalloidin: green; Alexa Fluor 488 and GLI1: Red (Alexa Fluor 647) and DAPI (Blue) staining (means ±SD, n=10; values by 2-tailed unpaired t-test P<0.05, **P<0.001, ****P<0.0001; images are representative of 3 individual biological replicates). (B) Quantification of propidium iodide-FACS staining of ONS76 cells after GYS32661 treatment at different time points. (C) Immunoblot representative images showing Cyclin D1 expression and its quantitation as compared to loading control (β Actin) in ONS76 cells after GYS32661 treatment at different time points (means ±SD, n=3; values by 2-tailed Student’s t-Test P<0.05, **P<0.001, ****P<0.0001) (Source data file).

**Supplementary Figure S7. GYS32661 treatment affects cell proliferation and intraciliary signaling in Shh-MB cells *in vitro***. (A) Representative images of ONS76 cells show cilia labeled with acetylated tubulin (red) and pericentrium (green) after GYS32661 treatment for 24 hours. Quantitative analysis was performed of cilia and pericentrium (means ±SD, n=3 different areas; values by 2-tailed Student’s t-Test P<0.05, **P<0.001, ****P<0.0001 representative of 3 individual biological replicates); (B-C) Representative immunoblots images show Ift88, pGsk3β, Total Gskβ, pVav2, Vav2, pPak1 and Pak1 levels with corresponding loading control (β Actin or GAPDH) after treatment with varying concentrations of GYS32661 for 24 hours in MSC4 cells. (D) Representative immunoblots show Acetylated tubulin (AcTub) and ARL13B expression with corresponding loading control β-Actin after GYS32661 treatment (Source data file).

## Notes

### Competing Interest Statement

The authors have declared no competing interest.

### Summary of Updates

The order of author names has been revised.

