## Supplementary Figures for "RAC1 Regulates Shh-Medulloblastoma Growth via GLI-Mediated Transcription"

Supplementry Figure S1

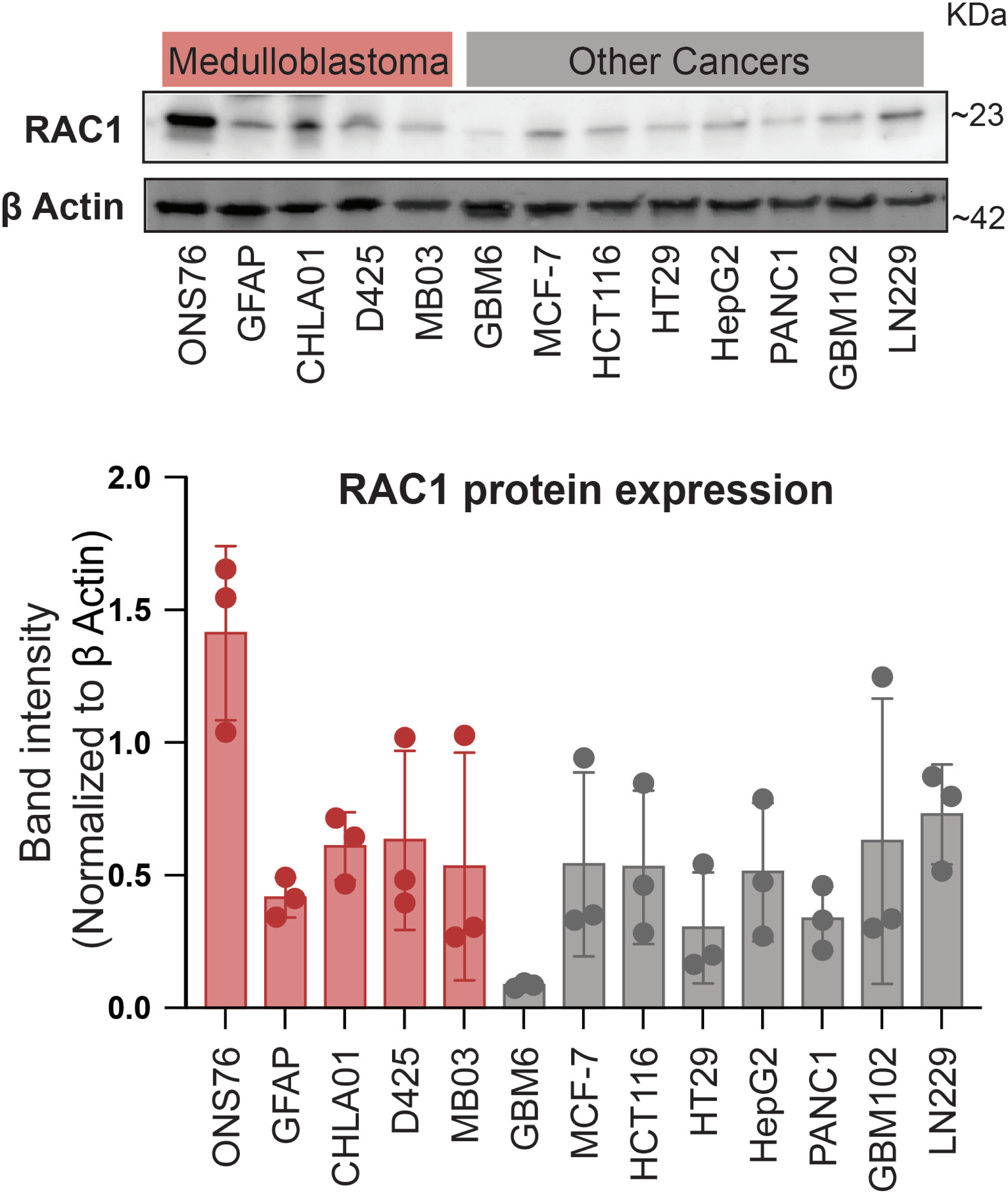

Supplementary Figure S2

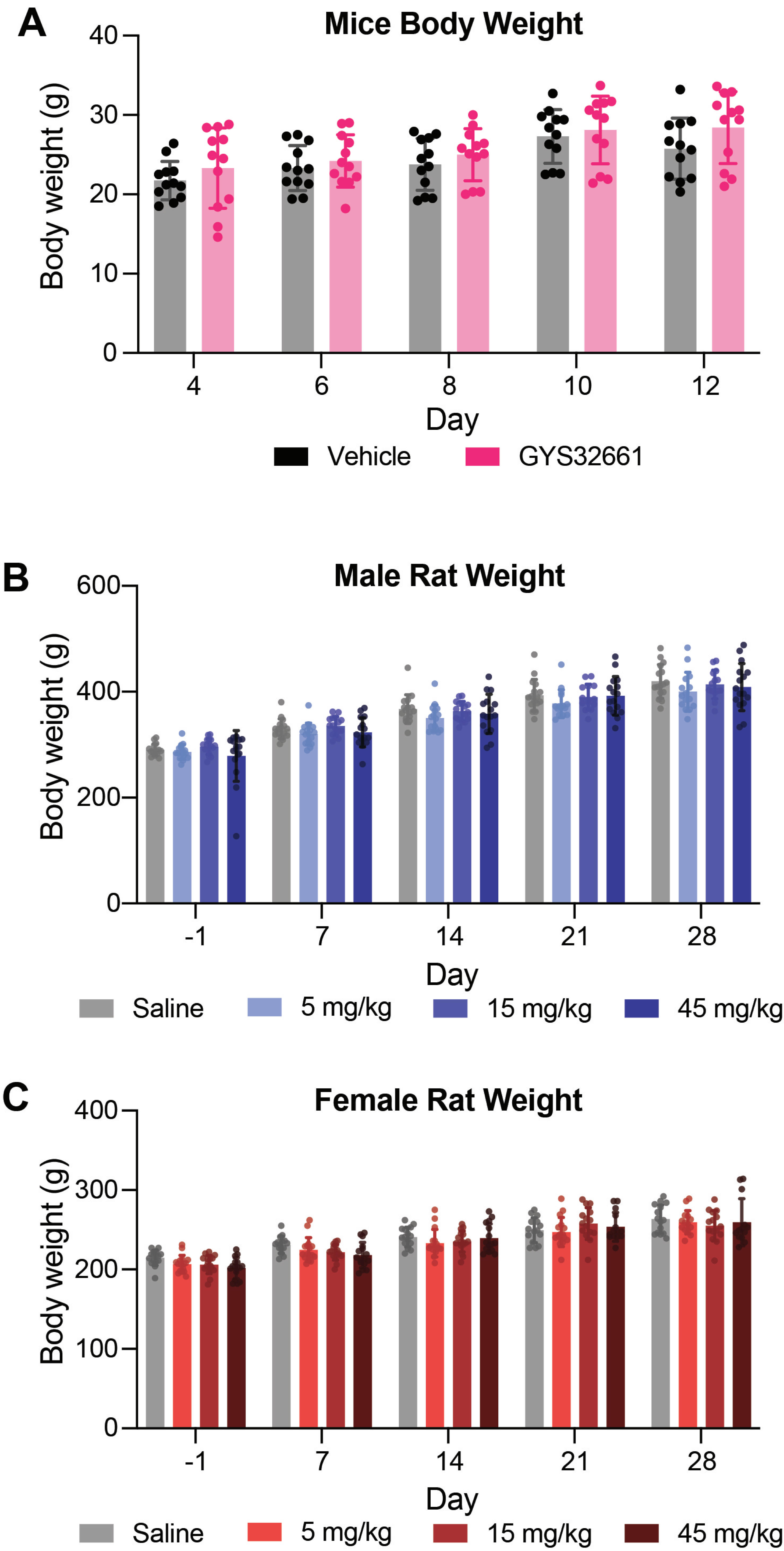

Supplementary Figure S3

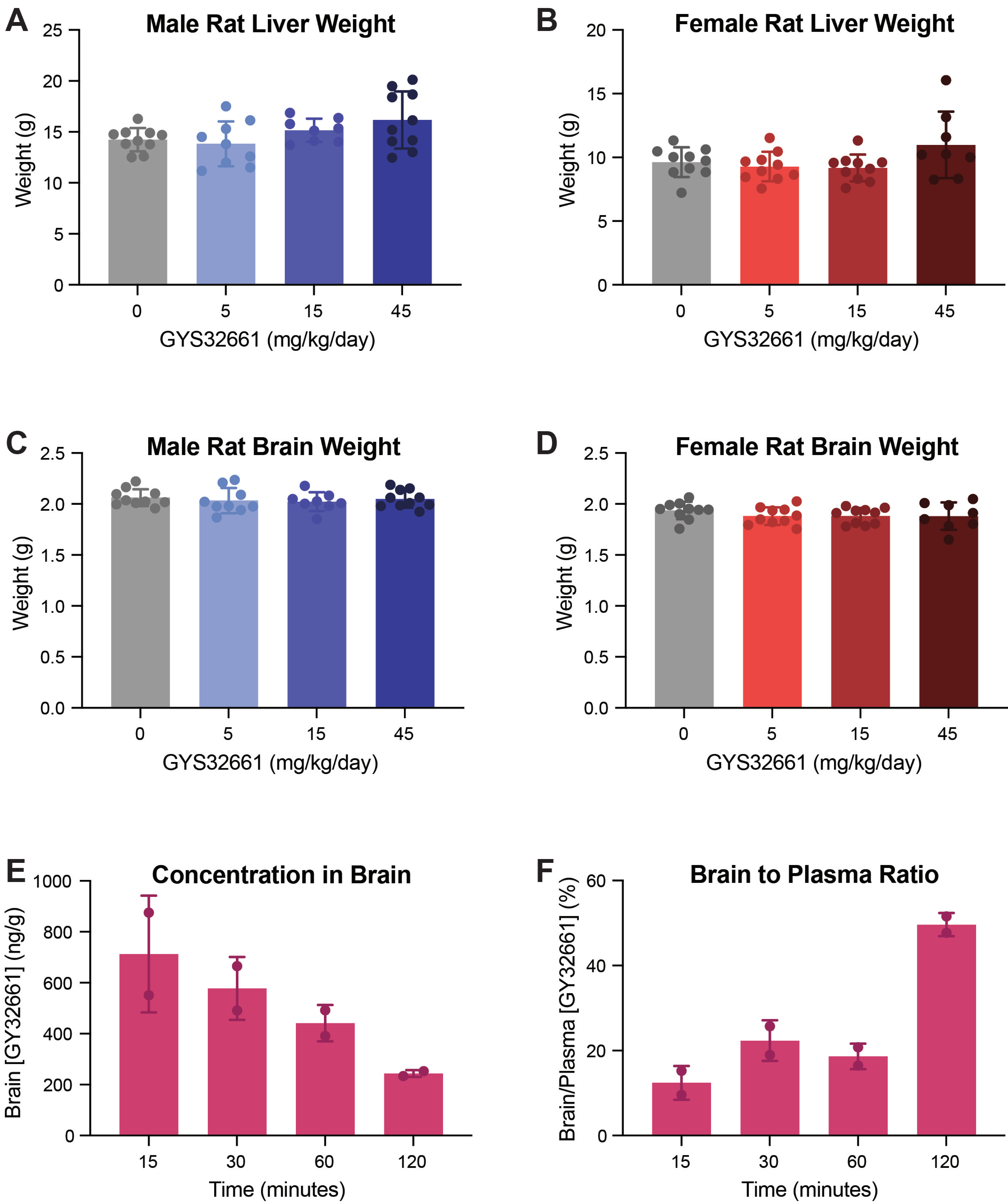

Supplementry Figure S4

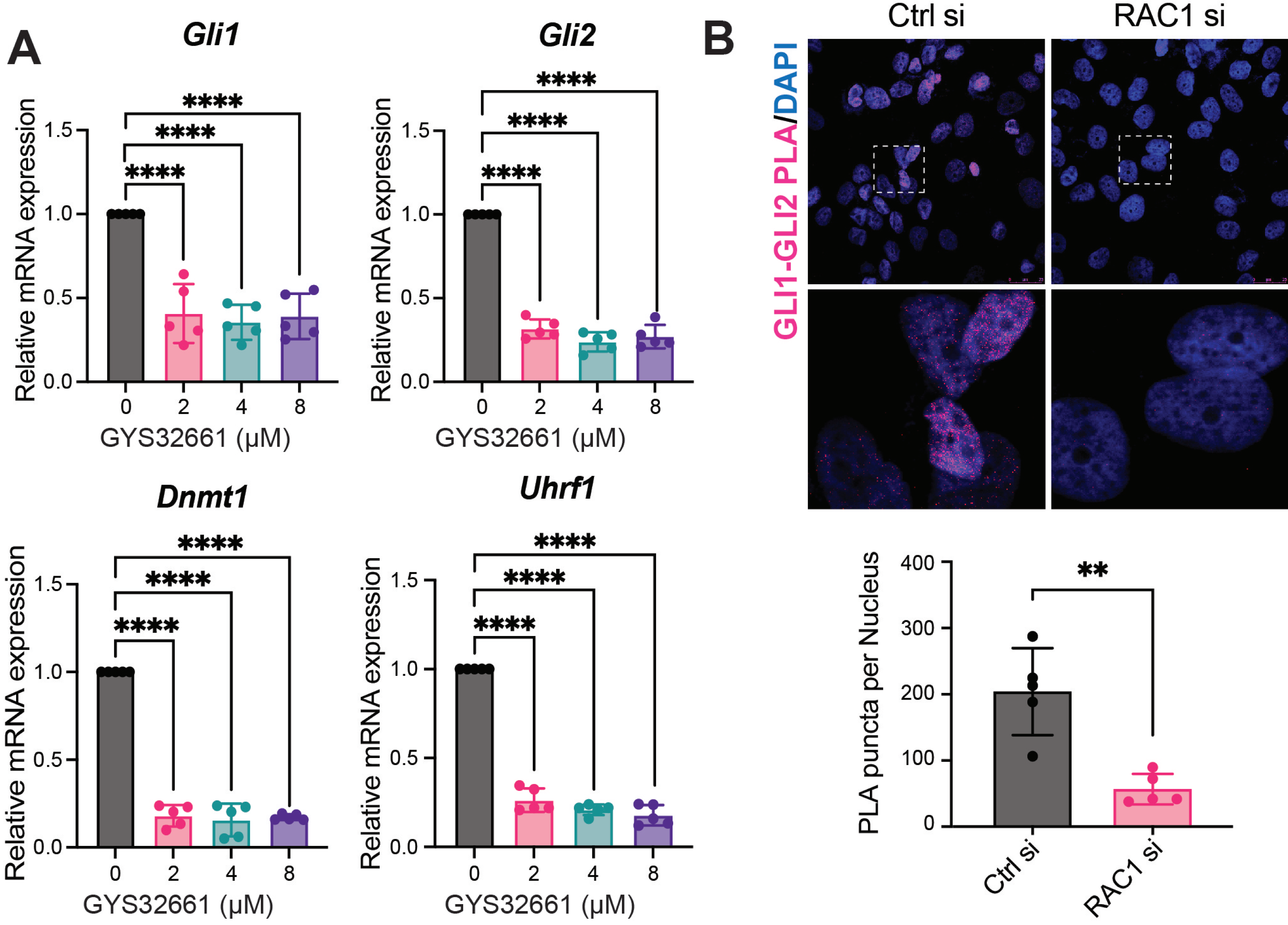

Supplementry Figure S5

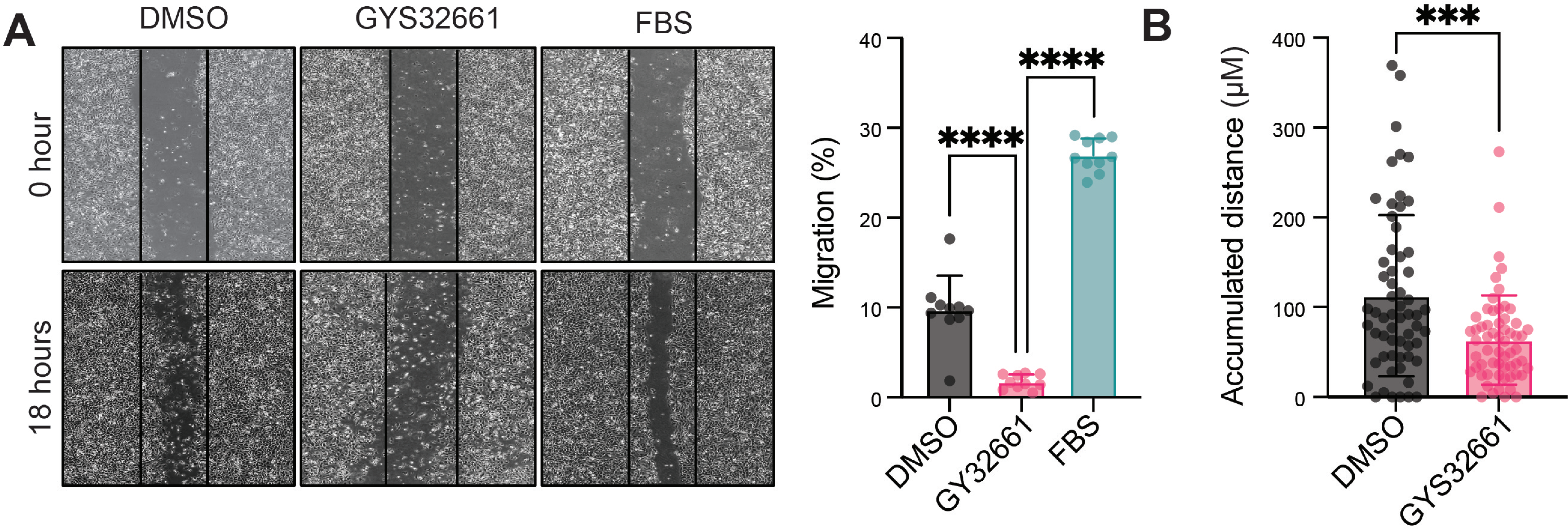

Supplementary Figure S6

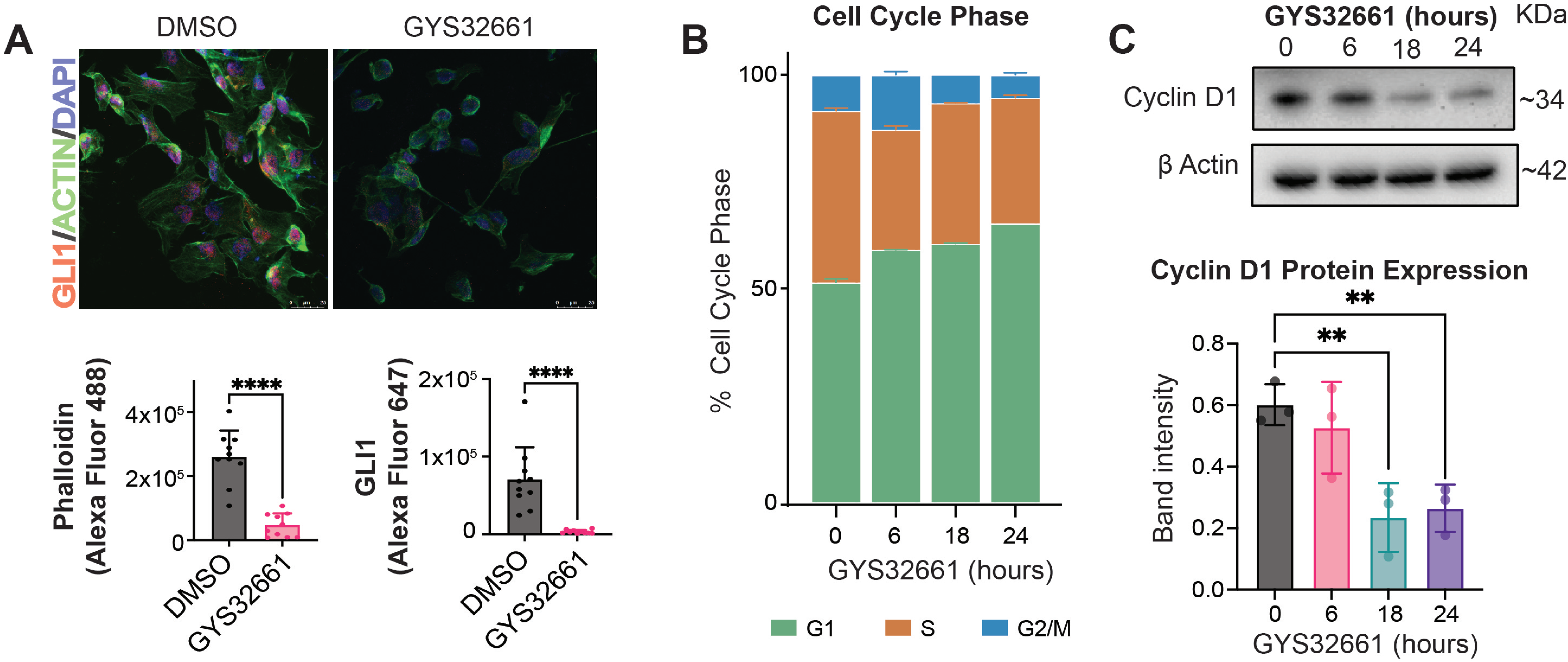

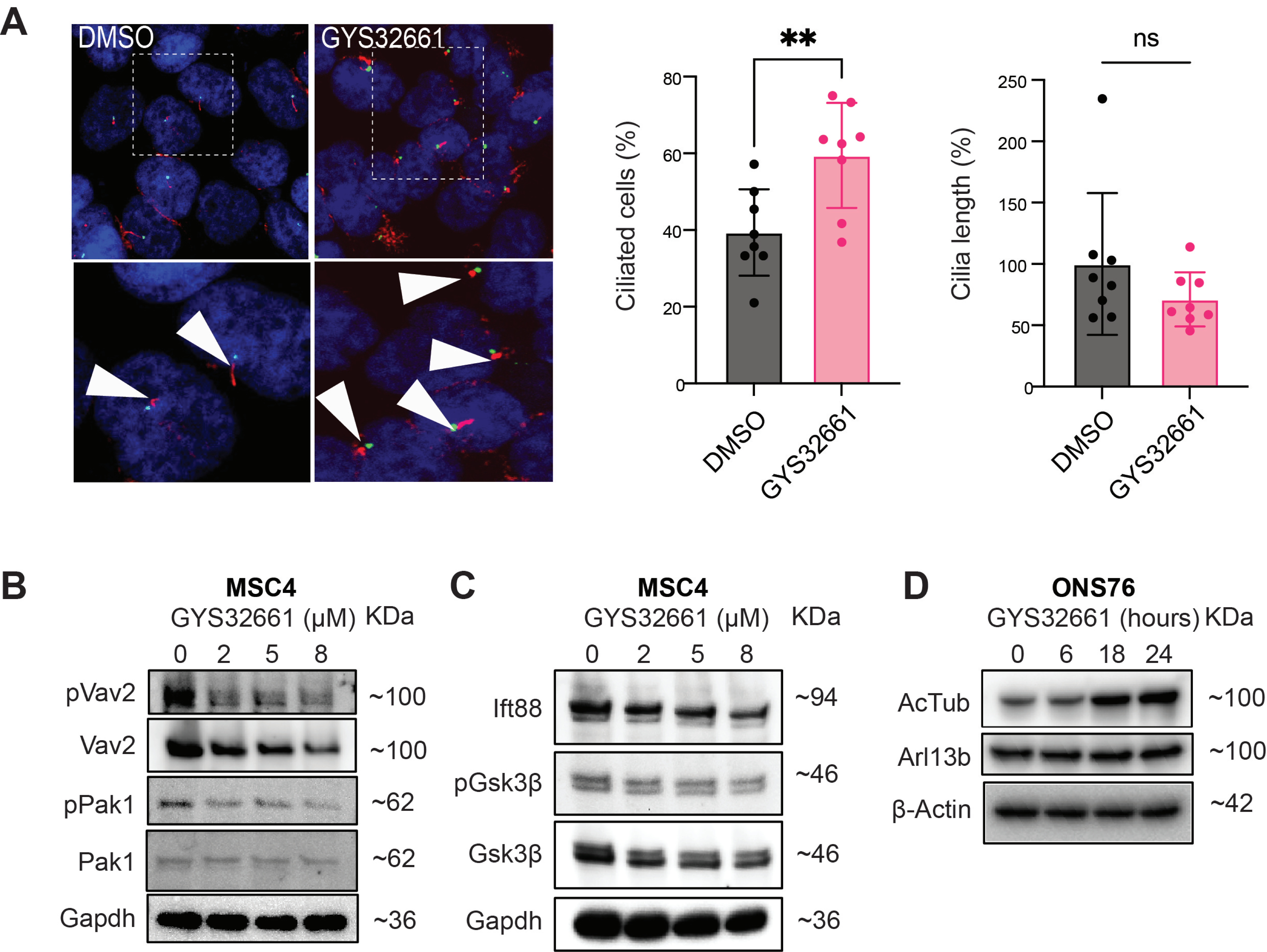
A
DMSO
GYS32661
**
ns
Ciliated cells (%)
0
20
40
60
80
DMSO
GYS32661
Cilia length (%)
0
50
100
150
200
250
DMSO
GYS32661
B
MSC4
GYS32661 (μM)
KDa
0 2 5 8
pVav2 ~100
Vav2 ~100
pPak1 ~62
Pak1 ~62
Gapdh ~36
C
MSC4
GYS32661 (μM)
KDa
0 2 5 8
Ift88 ~94
pGsk3β ~46
Gsk3β ~46
Gapdh ~36
D
ONS76
GYS32661 (hours)
KDa
0 6 18 24
AcTub ~100
Arl13b ~100
β-Actin ~42
