## Supplementary Table S1 for "RAC1 Regulates Shh-Medulloblastoma Growth via GLI-Mediated Transcription"

Supplementary Table S1: List of qRT-PCR primers

| Gene Name | Sequence |
|---|---|
| Human qPCR primers | |
| h_GLI1_F | CTACATCAACTCCGGCCAAT |
| h_GLI1_R | CGGCTGACAGTATAGGCAGA |
| h_GLI2_F | TGGAGCACTACCTCCGTTCT |
| h_GLI2_R | ACGAGGGTCATCTGGTGGTA |
| h_GAPDH_F | GTCAGTGGTGGACCTGACCT |
| h_GAPDH_R | TGCTGTAGCCAAATTCGTTG |
| h_DNMT1_F | AGTCCGATGGAGAGGCTAAG |
| h_DNMT1_R | TCCTGAGGTTTCCGTTTGGC |
| h_Rac1_F | CGCCCCCTATCCTATCCGCA |
| h_Rac1_R | GAACACATCGGCAATCGGCTTGT |
| h_UHRF1_F | CTGGTACGACGCGGAGAT |
| h_UHRF1_R | CGACAGTCGTTCAGAGAATCA |
| Mouse qPCR primers | |
| Gli1F | CTGGAGAGAGAGGAGAAG |
| Gli1R | TATGCTCACTGTTGATGT |
| Gli2F | CCAATGAGAAACCCTACATCTG |
| Gli2R | TTCACATGCTTGCGGAGT |
| Dnmt1F | CCTAGTTCCGTGGCTACGAGGAGAA |
| Dnmt1R | TCTCTCTCCTCTGCAGCCGACTCA |
| Uhrf1F | GCATCTACAAGGTGGTGAAG |
| Uhrf1R | AGGCTCTGTGTCATCTCGTC |
| 18SF | GCCCTGTAATTGGAATGAGTCCACTT |
| 18SR | CTCCCCAAGATCCAACTACGAGCTTT |
| Rac1F | GAAGCTGACTCCCATCACCTACCC |
| Rac1R | GGGGACAGAGAACCGCTCGGATAG |
