## Supplementary Table S2 for "RAC1 Regulates Shh-Medulloblastoma Growth via GLI-Mediated Transcription"

Supplementary Table S2: Human Chip qPCR primers for GLI1 upstream region (Accession No. NG_029564)

| Primer | Sequence | Upstream region (bp) |
|---|---|---|
| GLI1Loc2HF | CTAGGGAAAGGGGCTTCAGT | 4230-4250 |
| GLI1Loc2HR | CACCCTTTGGATGGAACTTG | 4404-4424 |
